# A simple coloring method to distinguish colonies of the yeasts *Lachancea thermotolerans* and *Saccharomyces cerevisiae* on agar media

**DOI:** 10.1101/2021.03.15.434753

**Authors:** Chrats Melkonian, Auke Haver, Marijke Wagner, Zakaria Kalmoua, Anna-Sophia Hellmuth, Sebastián N. Mendoza, Douwe Molenaar

## Abstract

The yeast *Lachancea thermotolerans* converts consumed sugar partly to lactic acid instead of ethanol and is therefore used together with *Saccharomyces cerevisiae* to produce wines with a lower alcohol content. Being able to distinguish these yeasts is important for quality control and quantitative assessment of the contributions of both yeasts to wine fermentations. Commonly used methods to routinely distinguish these organisms are indirect or rely on commercial products of undisclosed composition. Here we describe that adding bromocresol purple to agar media induces *Lachancea* colonies to develop a brown color, whereas *Saccharomyces* colonies remain white.

## 1. Introduction

*Lachancea thermotolerans* can be routinely distinguished from *Saccharomyces cerevisiae* by using selective media exploiting the property of *Saccharomyces* that it, in contrast to many other yeasts, can not use lysine as a sole nitrogen source [1]. However, using this selective medium, the proportions of *Saccharomyces* and *Lachancea* cells in mixed cultures can only be deduced by both counting colonies on selective and non-selective media. Therefore, this method is inherently less accurate than a method in which these yeast species can be distinguished on a single medium. In principle this is achieved by using “Chromagar Candida” (CHROMagar, Paris). On this commercially available medium which contains a chromogenic mix of undisclosed composition, colonies of both species are distinctly colored [1]. It has, however, not yet been demonstrated that this method can be routinely used to quantify both species in samples from mixed cultures.

In an attempt to find a simple way to distinguish these yeast colonies on a single plate we tried to use a bromocresol purple as a colorimetric pH indicator, which shifts from purple to yellow when the pH drops below 5.2. We assumed that the production of lactic acid would lead to a low pH near *Lachancea* colonies. However, also *Saccharomyces* colonies produce enough acid to decrease the pH, and usually the entire plate shifts color from purple to yellow. We report here the unexpected phenomena that after five days of incubation, however, *Lachancea* colonies develop a brown pigmentation, with increasing intensity from the edges to the center of the colony. In contrast, *Saccharomyces* colonies remain white, also after longer incubation (five days). This difference in color can be easily used to distinguish *Lachancea* and *Saccharomyces* colonies.

## 2. Results and Discussion

Rapid experimental techniques, such as flow cytometry and microscopy, are incapable to discriminate between *L. thermotolerans* and *S. cerevisiae* in mixed-culture, which indicates their high morphological similarity (not shown). Instead, using a medium with bromocresol purple shown positive results. After three to five days of growth, colonies of both yeasts are large and *L. theour-motolerans* colonies have a clear yellow first (day 3) and brown later (day 5) pigmentation (fig. 1). All *L. thermotolerans* colonies derived from four different strains develop the brown pigment, whereas all *S. cerevisiae* colonies remain white, as was checked by plating the species separately (not shown). Bromocresol purple is usually applied in growth media as a pH indicator. It has a pKa of 6.3 and changes color from purple to yellow when the pH drops from 6.8 to 5.2 [2]. Because of this property, bromocresol purple can be used to distinguish lactic acid producing bacteria in mixed cultures [3]. However, plates with either yeast species turned entirely yellow relatively soon, skipping a phase with distinct halos around colonies. Also, another commonly used pH indicator, methyl red, did not yield distinct halos around colonies of either species.

**Figure 1:**
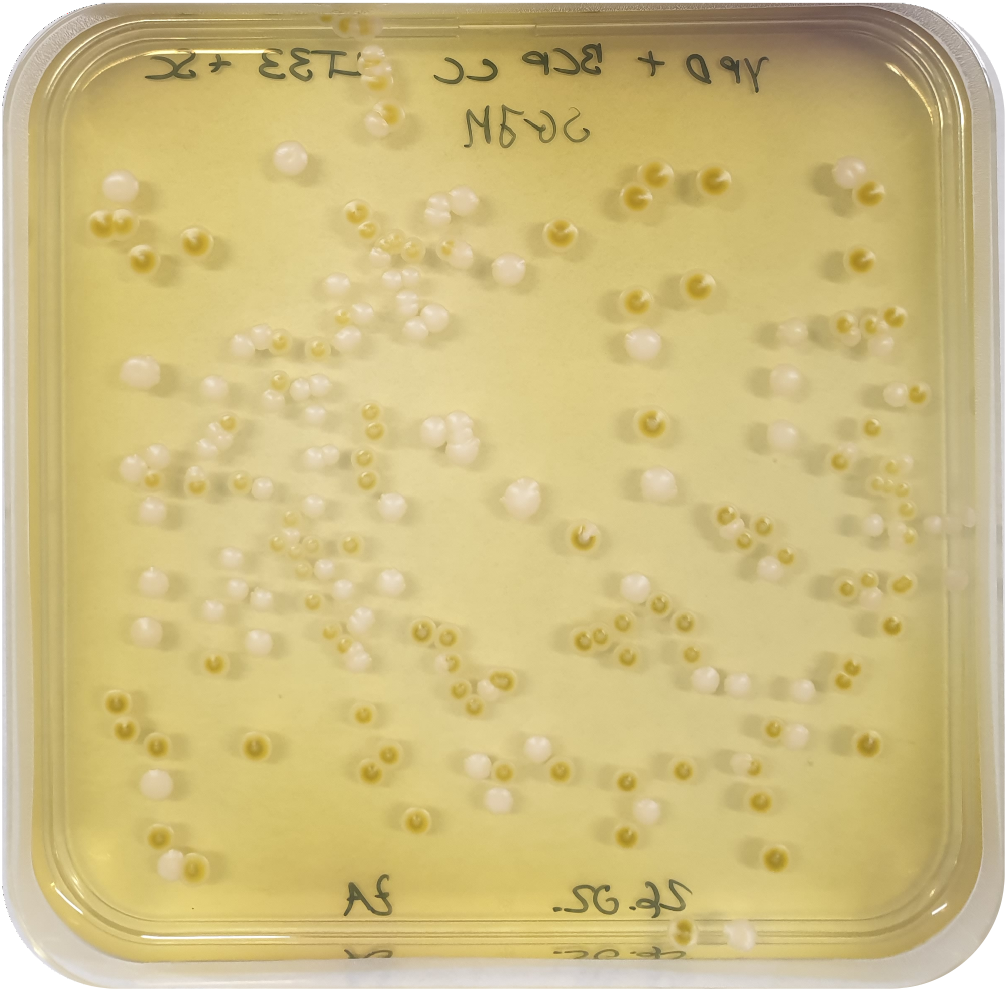
Colonies of *S. cerevisiae* and *L. thermotolerans* are distinguishable by their color on YPD plates containing bromocresol purple on which *L. thermotolerans* colonies develop a distinct brown pigmentation, which is most intense at the center of the colonies. The organisms were plated in a 1 to 1 ratio, and out of the four *L. thermotolerans* strains tested the one used here was NCYC 2433.

The cause of the brown pigmentation of *L. thermotolerans* colonies is unknown. The localized character of the staining, often more prominent in the center of the culture, suggests that the product that causes the staining does not diffuse freely. Therefore, likely mechanisms are an intracellular chemical conversion and containment or metachromasy, *i.e*. a colour change by binding specifically to a high molecular weight component of *Lachancea* cells [4, 5].

The method proposed here was tested for different ratios of mixtures of the yeasts, ranging from 1:30–30:1 and yielded results that were in reasonable agreement with the theoretical values (fig. 2). The maximal deviation out of 14 measurements was 2.7 fold from the theoretical value, and 9 out of 14 values deviated by less than 1.25-fold from the theoretical value. This demonstrates that this rapid quantification method for mixed cultures with *Lachancea* is useful for practical purposes.

**Figure 2:**
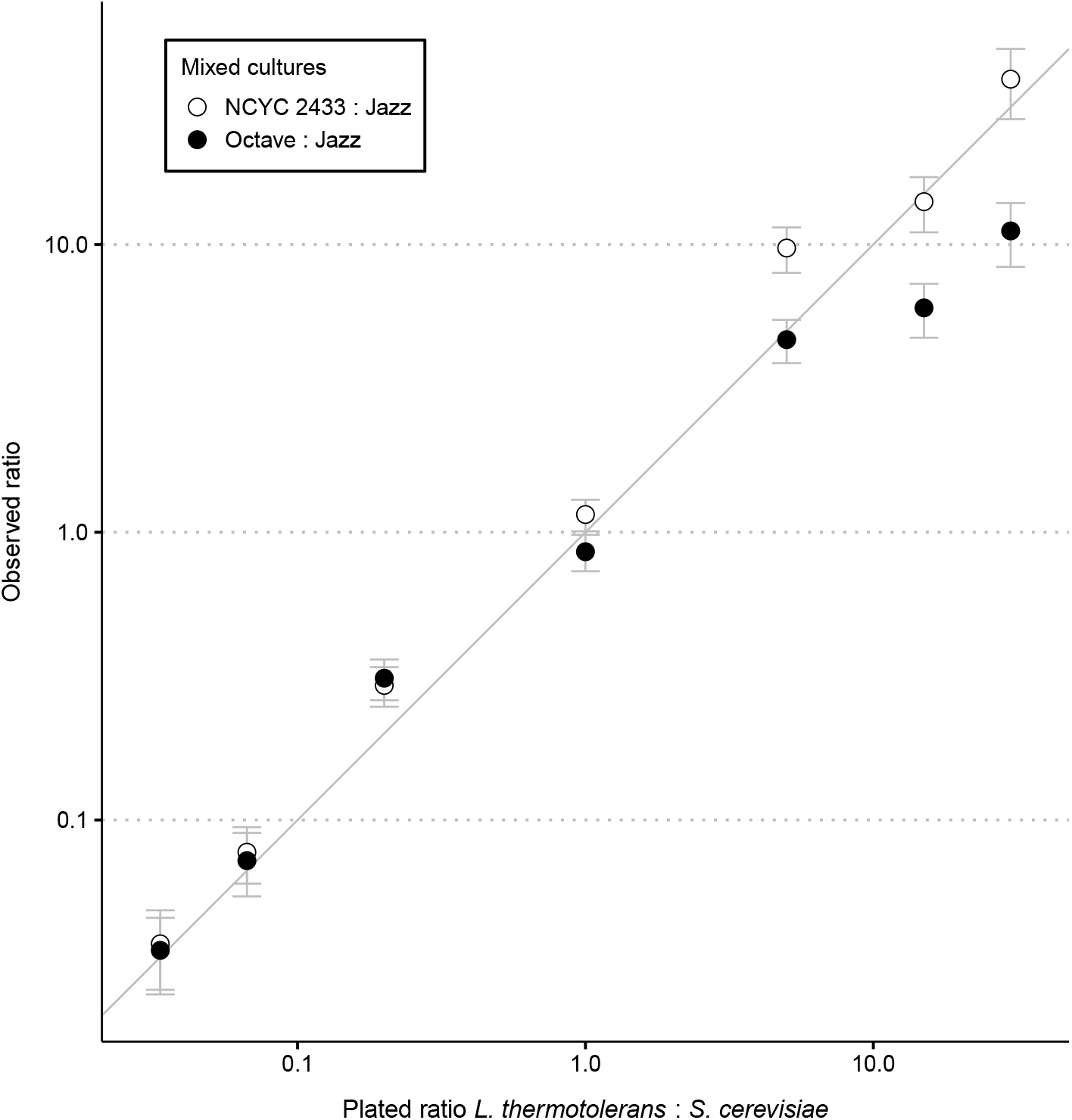
Comparison of plated ratio of two *L. thermotolerans* stains to *S. cerevisiae* and observed ratio of colonies. The gray line indicates the identical values of both variables and the bars indicate expected standard deviations assuming a Poisson distribution of the error of colony counts. Both axes have a logarithmic scale.

## 3. Materials and Methods

### 3.1. Yeast strains

*Lachances thermotolerans* strains NCYC 2433 (type strain), Viniflora^®^ Octave, Viniflora^®^ Concerto and a strain from a private collection (Chr. Hansen, Denmark) were used as well as *Saccharomyces cerevisiae* strain Viniflora^®^ Jazz (Chr. Hansen, Denmark).

### 3.2. Growth media

Synthetic grape juice medium (SGJM) consisted of D-glucose (110 g/l), D-fructose (110 g/l), L-tartaric acid (6 g/l), L-malic acid (3 g/l), citric acid (0.5 g/l), Yeast Nitrogen Base without amino acids and without ammonium sulfate (YNB, Sigma, 1.7 g/l), calcium chloride (0.2 g/l), casein hydrolysate (0.2 g/l), arginine monohydrochloride (0.8 g/l), L-proline (1 g/l), L-tryptophan (0.1 g/l) and inositol (0.8 g/l). To produce the SGJM, three solutions (A, B and C) are made and mixed together, solution A consists of D-glucose and D-fructose in 450 ml of demineralized water, solution B of L-tartaric acid, L-malic acid and citric acid dissolved in 200 ml of demineralized water, and solution C of YNB, calcium chloride, casein hydrolysate, arginine monohydrochloride, L-proline, L-tryptophan and inositol dissolved in 200 ml of demineralized water. After adding all solutions together, the pH was adjusted to 3.5 using 5 M KOH, and demineralized water was added to a final volume of 1 L. The medium was filter-sterilized [6].

Yeast extract Peptone Dextrose agar (YPD-agar) consisted of yeast extract (2 g/l), peptone (4 g/l), agar (15 g/l). When desired, 25 mg/l of bromocresol purple was added to this medium. The pH was adjusted to between 6.8 and 7 using 5 M KOH, and the medium was sterilized by autoclaving, upon which 40 ml of filter-sterilized glucose (550 g/l) was added. Negative controls were performed with the addition of 80 mg/l of methyl red instead of bromocresol purple.

### 3.3. Microscopy and flow cytometety

Images were acquired using an Olympus BX63 wide-field epi-fluorescence microscope with a 100X/1.35NA UPlanApo objective. Samples were visualized using an LED Lumencore SOLA SE FISH and the ORCA-Fusion Digital CMOS camera (Hamamatsu). Image pixel size: XY, 64.5 nm. CellSens software (Olympus) was used for acquisition and Differential Interference Contrast (DIC) was applied. For flow cytometry measurements (forward and side scatter as well as green fluorescence) an Accuri C6 flow cytometer (BD Biosciences, San Jose, CA, US) with the corresponding software, CFlow Plus, was used.

## Acknowledgements

We thank Laura Luzia, Evelina Tutucci, Emile Zwering, Lorenzo Peyer and Bas Teusink for their significant contribution with discussions. We thanks Chr. Hansen for providing the strains. The research was partially supported by a Grand Solution grant from Innovation Fund Denmark (grant no. 6150-00033B), The FoodTranscriptomics project.

## Author contributions Statement

C.M. and D.M. conceived the methodology. A.H., Z.K and A.S.H performed the experimental work supervised by M.W., S.N.M and C.M. D.M. and C.M. wrote the manuscript.

